# Single-cell morphological tracking of liver cell states to identify small-molecule modulators of liver differentiation

**DOI:** 10.1101/2023.11.15.567184

**Authors:** Rebecca E. Graham, Runshi Zheng, Jesko Wagner, Asier Unciti-Broceta, David C. Hay, Stuart J. Forbes, Victoria L. Gadd, Neil O. Carragher

## Abstract

Alternative therapeutic strategies are urgently required to treat liver disease, which is responsible for 2 million deaths anually. By combining Cell Painting, a morphological profiling assay that captures diverse cellular states, with the bi-potent HepaRG^®^ liver progenitor cell line, we have developed a high-throughput, single-cell technique, to track liver cell fate and map small-molecule induced changes using a morphological atlas of bi-lineage liver cell differentiation. To our knowledge this is the first-time single-cell trajectory inference has been applied to image-based Cell Painting data and leveraged for drug screening. The overarching goal of this new method is to aid research into understanding liver cell regeneration mechanisms and facilitate the development of cell-based and small-molecule therapies. Using this approach, we have identified a class of small-molecule SRC family kinase inhibitors that promote differentiation of HepaRG^®^ single-cells towards the hepatocyte-like lineage and promotes differentiation of primary human hepatic progenitor cells towards a hepatocyte-like phenotype *in vitro*.

## Introduction

Hepatocytes are long-lived quiescent cells that, within the context of the liver *in vivo,* retain their proliferative capacity, thus allowing for homeostatic and injury-induced liver regeneration ^1,2^. However, in chronic liver disease the proliferative capacity of hepatocytes is diminished. The expansion of hepatic progenitor cells, which have the potential to differentiate into functional hepatocytes, represents an alternative pathway for liver regeneration ^3,4^. However, the extent of the contribution and the mechanisms responsible for this alternative pathway are unclear, and the only curative option for patients with severe chronic liver disease is liver transplantation. Unfortunately, donor organ availability does not meet demand, and, for example, in the UK approximately 20% of patients die waiting for a donor ^5^, making liver disease responsible for 4% of all deaths (1 in every 25 deaths) worldwide accounting for two million deaths annually ^6^. Therefore, novel therapeutic strategies are urgently required for the treatment of advanced liver disease.

Both cell-based and small-molecule therapies hold promise as alternative strategies to whole organ transplant ^7,8^. However, two major barriers remain; firstly, *in vitro* proliferation of adult human hepatocytes is limited and challenging ^9,10^ and the generation of functionally mature liver cell types *in vitro* is only partial since primary human hepatocytes rapidly lose most of their differentiated functions ^11^. These challenges limit the application of cell-based therapies and the study of liver cell functions *in vitro* and must be overcome for the liver research field to exploit alternative therapeutic strategies to treat liver disease. This project therefore aims to identify small-molecule modulators of liver progenitor cell status, particularly those that promote the generation of fully mature liver cell types, to aid cell banking, research into liver cell regeneration mechanisms, and the development of cell-based and small-molecule therapies to treat liver disease.

To track cell state transitions involved in liver cell differentiation we sought to use multiparametric high-content cellular morphological profiling which offers an inexpensive, high-throughput, single-cell, spatially resolved alternative to RNA-sequencing for profiling cell states. High-content imaging is having a major impact at all stages of the drug discovery process ^12^. However, despite collecting single-cell information within each image most data analysis is performed on aggregated cell populations at the “well level”, and single-cell analysis pipelines have not been developed ^12,13^. In 2013 the Cell Painting assay was developed as an unbiased, cell based, morphological high-content profiling assay capable of capturing subtle changes in morphology ^14^. It combines six fluorescent dyes, imaged in five channels, to visualise eight cellular components and organelles; nuclei, F-actin, endoplasmic reticulum, mitochondria, golgi, plasma membrane, cytoplasmic RNA and nucleoli ^14–16^. Quantitative data can then be extracted from microscopy images to identify the phenotypic impact of chemical or genetic perturbations. Due to the broad morphological information contained in the images, Cell Painting has been exploited for many research applications, including clustering chemical and genetic perturbations based on their morphological impact, identifying disease phenotypes, drug screening, toxicity prediction and mechanism of action prediction using machine learning ^17–23^.

By combining Cell Painting with the bi-potent HepaRG^®^ liver progenitor cell line, we have developed a high-throughput single-cell technique to track liver cell fate during bi-lineage differentiation. In this work we describe the first morphological assay to leverage the mathematics of trajectory inference and apply it to high dimensional, single-cell, morphological data to track liver cell fate and identify small-molecule modulators of liver cell differentiation.

The HepaRG^®^ cell line is a popular *in vitro* model and primary human hepatocyte surrogate ^24^. The HepaRG^®^ progenitor cells demonstrate bi-lineage differentiation potential and produce a mixed population of hepatocyte-like and biliary-like cells over the course of four weeks with the addition of commercially available differentiation supplements halfway through the process ^25^. Using the Cell Painting assay to capture the morphological changes induced by a bioactivity compound library of 496 target-annotated small molecules we identified compounds that induced the same differentiated morphological phenotype as the differentiation supplement at the image level. We then developed a single-cell analysis pipeline to track liver cell differentiation using the Cell Painting data and applied trajectory inference to build a single-cell morphological atlas of liver cell differentiation. Finally, we overlaid the drug induced data to gain single-cell resolution into the drug induced changes to assess how the compounds altered liver cell fates and lineages.

From this work we have identified compounds and their putative targets and pathways that support expansion and appropriate lineage commitment, to aid the development of cell-based and small-molecule therapies to treat liver disease. Further, we have taken our hepatocyte promoting compounds, including SRC family kinase (SFK) inhibitors and validated them in primary human hepatic progenitor cells, where partial differentiation into hepatocytes is inferred through an increase in hepatic functional markers such as HNF4a and Cyp2E1.

Overall, we have developed a high-throughput morphological cell state tracking pipeline at the single-cell level to identify small molecule modulators of liver cell differentiation which were subsequently validated in lower throughput primary human hepatic progenitor differentiation assays. Single-cell morphological profiling represents a holistic, high-throughput and inexpensive method to quantify cell state transitions and is applicable to many assays systems, providing deeper insights into the impact of small-molecule perturbations on cellular differentiation.

## Results

### Cell Painting can be used to morphologically distinguish liver cell states

The bi-potent liver progenitor cells HepaRG^®^, undergo differentiation producing a mixed population of biliary-like and hepatocyte-like cells. During this differentiation process the cells undergo distinct morphological changes (**Fig. 1a**). To identify molecules that modulate liver cell differentiation, we developed a high-content morphological assay protocol in 384-well plate format, compatible with automated high-throughput imaging, to track liver cell differentiation by applying Cell Painting to the bi-potent HepaRG^®^ cells (**Fig. 1b**).

**Fig 1.**
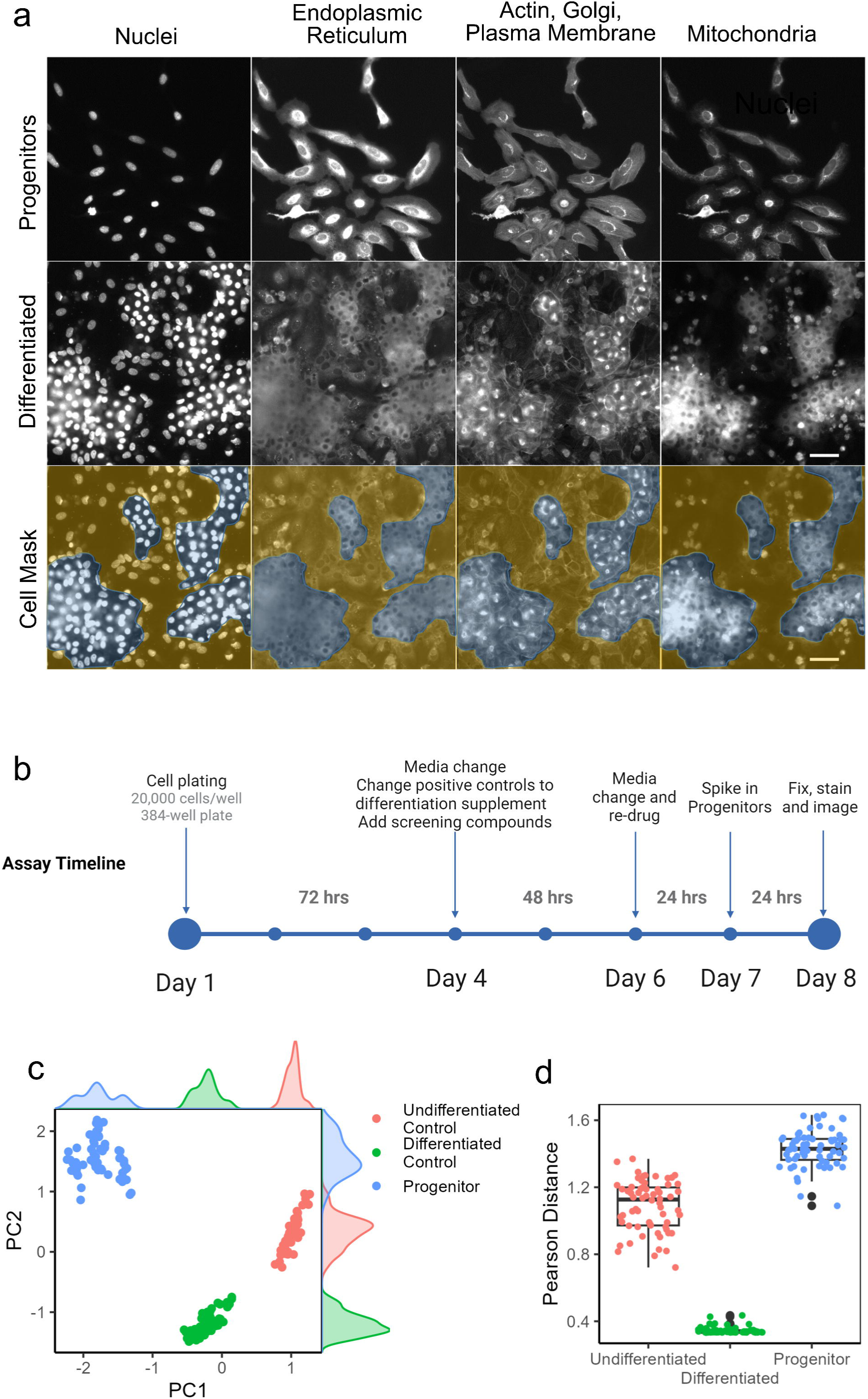
High-content Cell Painting assay in HepaRG^®^ cells. a) Cell Painting images of nuclei (Hoechst), endoplasmic reticulum (Concanavalin A), F-actin, golgi and plasma membrane (Phalloidin and Wheatgerm agglutinin), and mitochondria (MitoTracker DeepRed) across HepaRG^®^ progenitors and HepaRG^®^ differentiated cells. Hepatocyte-like cells (blue mask) and cholangiocyte-like cells (yellow mask). b) Timeline for 8-day assay. c) Principal component analysis of assay controls at the image level morphology across 4 replicates. d) Pearson distance from differentiated controls across all control images.

To begin with we first assessed the ability of the Cell Painting assay to distinguish the HepaRG^®^ progenitors from our positive control, HepaRG^®^ Differentiation Medium Supplement, and negative control, HepaRG^®^ Growth Medium Supplement (here after referred to as differentiated control and undifferentiated control) after eight days in cell culture. Initial analysis was performed using the well level median morphology metrics, an industry standard approach for high-content drug screening. At the end of the eight days the differentiated controls are partially differentiated, forming a mix of hepatocyte-like and biliary-like cells (**Supplementary Fig. 1**). This assay format allows us to identify modulators of liver cell differentiation including those that increase differentiation of the cells beyond that of the differentiated controls within the 8-day assay period.

Quantification of the controls was carried out using the median cell value for 848 morphological features extracted using Cell Profiler. Principal component analysis and Pearson distance to the positive controls across four separate plate replicates shows clear separation of the controls and reproducible differentiation phenotypes (**Fig. 1c and d**). Coefficient of variation (CV) for Pearson distance ranges from 1.2-14.6 % across the control classes and plate Z-primes range from 0.17-0.65 (**Supplementary Fig. 2**) across four plates. Further, a three-class Random Forest machine learning classifier trained to distinguish the controls from each other performed with an accuracy score of 1 (100% accurate) across the three classes. Together this demonstrates a robust assay for drug screening.

### Identification of small-molecule modulators of liver cell bilineage differentiation

We used this assay to profile 496 compounds from a library of bioactive target annotated compounds (**Supplementary Table 1**) screened at two concentrations (5µM and 0.5 µM) to identify hit compounds that promote the differentiation of HepaRG^®^ liver cells. We identified 27 hits that had a Pearson distance of less than 0.9 to the differentiated control (+differentiation supplement) and a Euclidean distance of less than 1.7 (**Fig. 2a and b**). Hits belong to several classes of compounds including HDAC inhibitors, PI3K inhibitors, and SRC family kinase (SFK) inhibitors (**Supplementary Table 2**). HDAC inhibitors have previously been shown to promote the differentiation of induced pluripotent stem cells into hepatocyte-like cells ^26^, highlighting the ability of our HepaRG^®^ cell assay to identify biologically relevant small-molecules and pathways.

**Fig 2.**
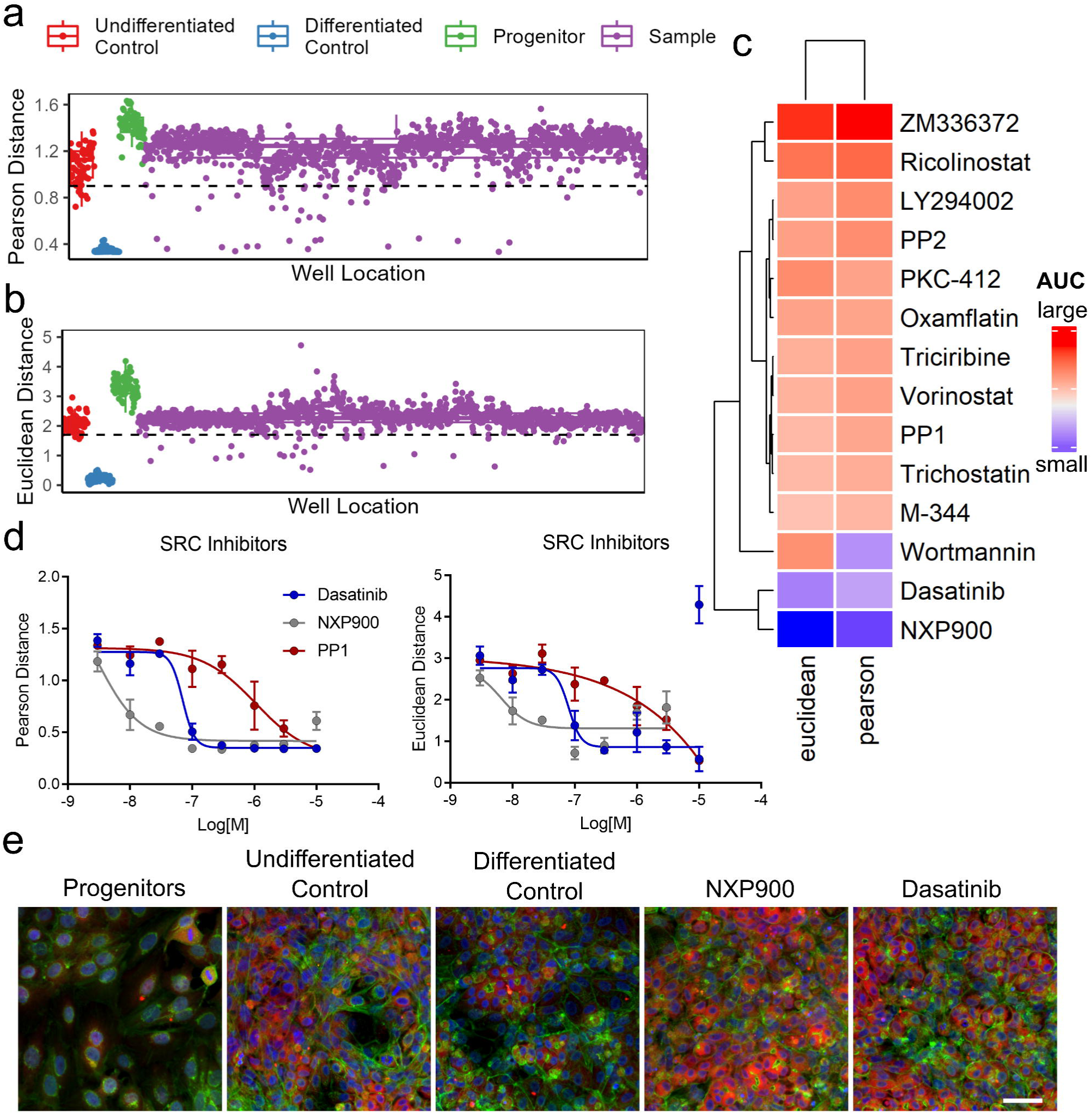
Drug screen analysis. a) Pearson distance and b) euclidean distance from differentiated controls (blue) across all controls and 496 drug treatments (purple) at 5 and 0.5µM. Hit thresholds denoted by dashed lines. c) Area under the curve (AUC) for multiparametric dose responses across 14 compounds of interest. d) Multiparamentric dose responses across SFK inhibitors for Pearson and Euclidean distance. n = 3. e) Colour combined images for controls and top two hits (NXP900 and Dasatinib). Blue = Hoechst (nuclei), Green = Phalloidin and Wheatgerm agglutinin (F-actin, golgi and plasma membrane), Red = Mitotracker DeepRed (Mitochondria). Scale bar =50 µm.

To confirm initial compound screening hits we performed 8-point semi-log dose response tests on resupplied compound material. Further, noting that PP1 and PP2 both belong to the pyrazolopyrimidine class of cell permeable compounds that has been shown to inhibit SFK members; Lck, Fyn, Hck, and Src, as well as non-Src family kinases such as CSK, RIP2, and CK1δ ^27^, we added two further SFK inhibitors (dasatinib; a dual SRC/ABL inhibitor ^28^ and NXP900; a highly selective SFK inhibitor ^29^) to the list of compounds in order to determine if PP1 and PP2 were acting via SFK inhibition or other kinase targets. In total we re-tested 14 compounds in triplicate multiparametric dose responses (**Supplementary Fig. 3**). All compounds validated showing dose dependent activity except ZM336372 (**Fig. 2c**). Overall, the most potent hits were the SFK inhibitors, all of which validated, with the most selective SFK inhibitor, NXP900, being the most potent hit with an IC50 of 3nM (**Fig. 2c, d and e**).

### Single-cell morphological analysis

To gain insights at the single-cell resolution and to understand how the compound hits modulate cell fate towards the two different cell lineages, we built a single-cell morphological atlas of HepaRG^®^ bilineage differentiation and then overlaid the single-cell drug induced changes on top.

To build the morphological atlas of HepaRG^®^ differentiation we captured Cell Painting images every 24-72 hrs from 20 time points over the course of the full four-week differentiation protocol to track the morphological changes associated with HepaRG^®^ bilineage differentiation at the single-cell level (**Fig. 3**). UMAP analysis of the single-cell time course data over the entire differentiation process generates a continuous trajectory of cellular bilineage differentiation from a starting population of progenitors (Day 1) all the way through to two distinct final cell populations (Day 28) (**Fig. 3b**).

**Fig. 3.**
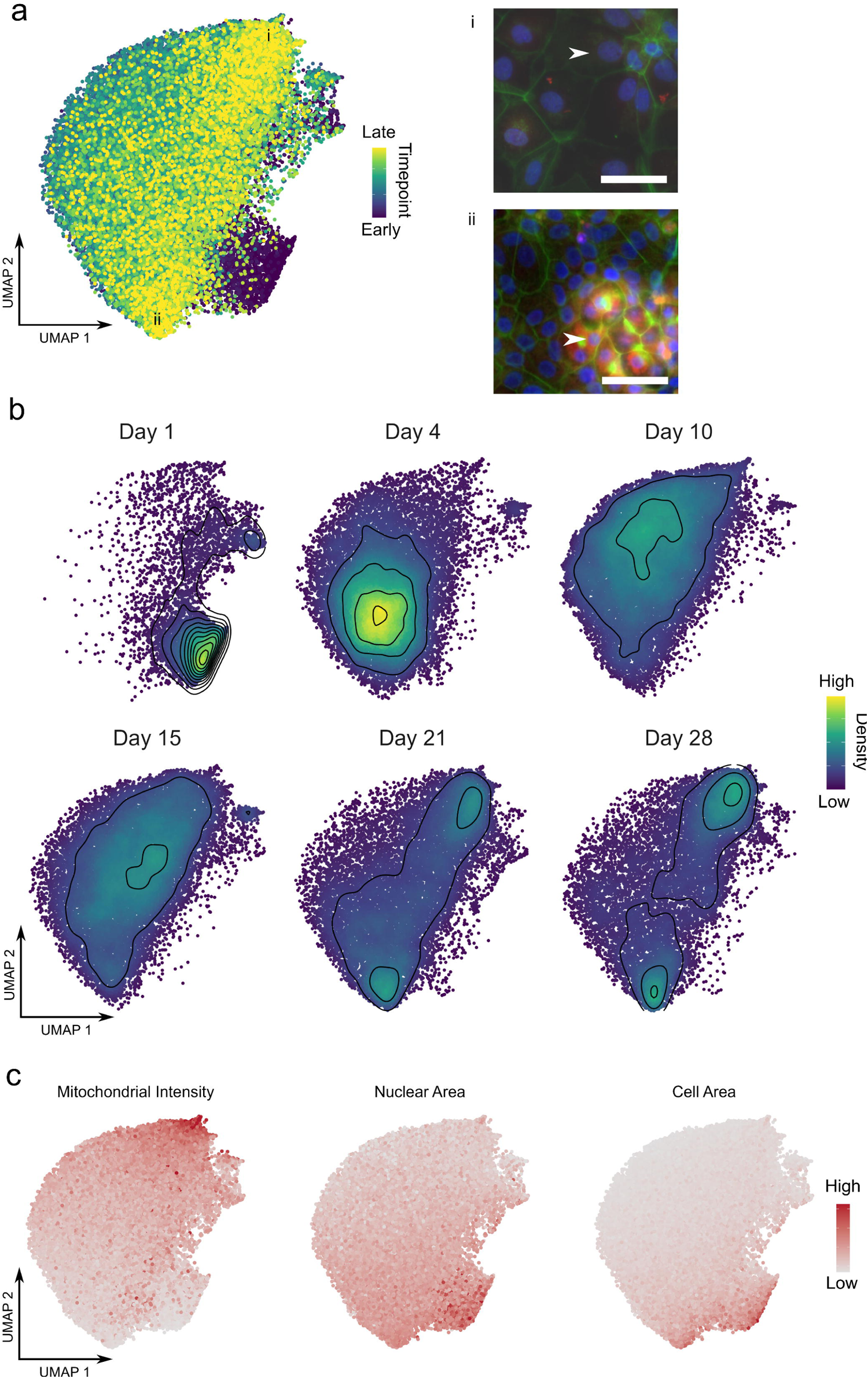
Single cell morphological atlas of HepaRG^®^ differentiation. a) UMAP embeddings of single-cell HepaRG^®^ timecourse Cell Painting data coloured by the timepoint at which the cell was imaged. Exemplar colour-combined images of single-cells (white arrows) belonging to the two clusters of differentiated, (final timepoint) cells (i and ii). Blue = Hoechst (nuclei), Green = Phalloidin and Wheatgerm agglutinin (F-actin, golgi and plasma membrane), Red = Mitotracker DeepRed (Mitochondria). Scale bar =50 µm. b) UMAP embeddings separated out across 6 timepoints spanning the differentiation period. Points coloured by density on plot and density contours added. C) UMAP embedding coloured by characteristic morphological features delineating the hepatocyte-like cells, biliary-like cells and progenitors.

To confirm that the two final cell populations are the expected clusters of hepatocyte-like and biliary-like cells we used known cellular morphological markers to identify the cell populations, similar to the use of gene expression markers in single-cell RNA-seq analysis. Here we used mitochondrial intensity to identify the metabolically active mature hepatocyte-like population, and cell and nuclear size to identify the biliary-like and progenitor populations (**Fig. 3c**). Overall the divergence of two separate cell populations demarked by expected morphological features is clear, demonstrating the ability of Cell Painting to track a complex process such as bilineage differentiation at the single-cell level.

Having confirmed the ability of the assay to resolve the different cell populations and build a continuous morphological atlas of the bilineage differentiation, we next overlaid the small molecule compound screening data. Firstly, we assessed the effects of the controls at the single-cell level (**Fig. 4a**). Importantly, the “spiked in” progenitor control from the screen (seeded 24hrs before the end of the assay) overlay completely with the original 24hr timepoint from the timecourse data used to create the single-cell morphological atlas (**Supplementary Fig. 4a**), demonstrating the ability to merge datasets at the single-cell level after plate-based standardisation to the controls. Secondly, the differentiated control samples in the screening assay push the cells closer to the final timepoints in the timecourse data compared to the undifferentiated control but the cells do not reach full maturity (**Fig. 4a**). This was expected since the screening assay was only 8 days long.

**Fig. 4.**
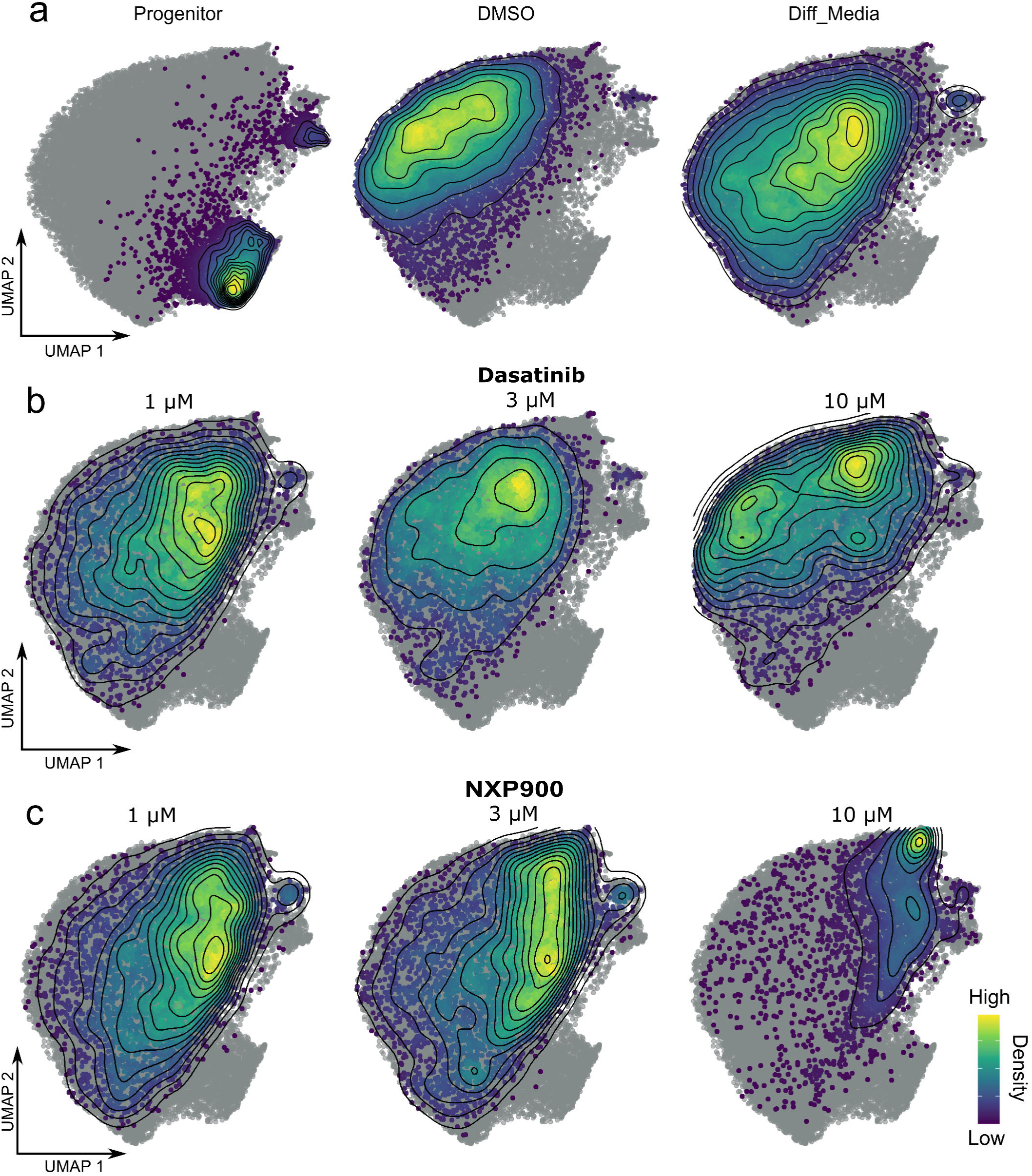
Single-cell drug treatments mapped to HeaRG atlas. a) Screening controls, b) Dasatinib and c) NXP900 single-cell data coloured by density and overlaid on the HepaRG^®^ atlas (grey point).

We next overlaid the single-cell data for compound treatments of interest at three concentrations (1, 3 and 10 µM). The confirmed compound treatments show dose dependent shifts in the cell populations (**Fig. 5b and c and Supplementary Fig. 5**). Consistent with the image level analysis, the SFK inhibitors demonstrate the greatest effects (**Fig. 4b and c**) and again, the selective SFK inhibitor NXP900 was found to be the most potent. Of interest both the approved SFK inhibitor, Dasatinib and particularly NXP900, very clearly favour the hepatic lineage in a dose dependent manner.

**Fig. 5.**
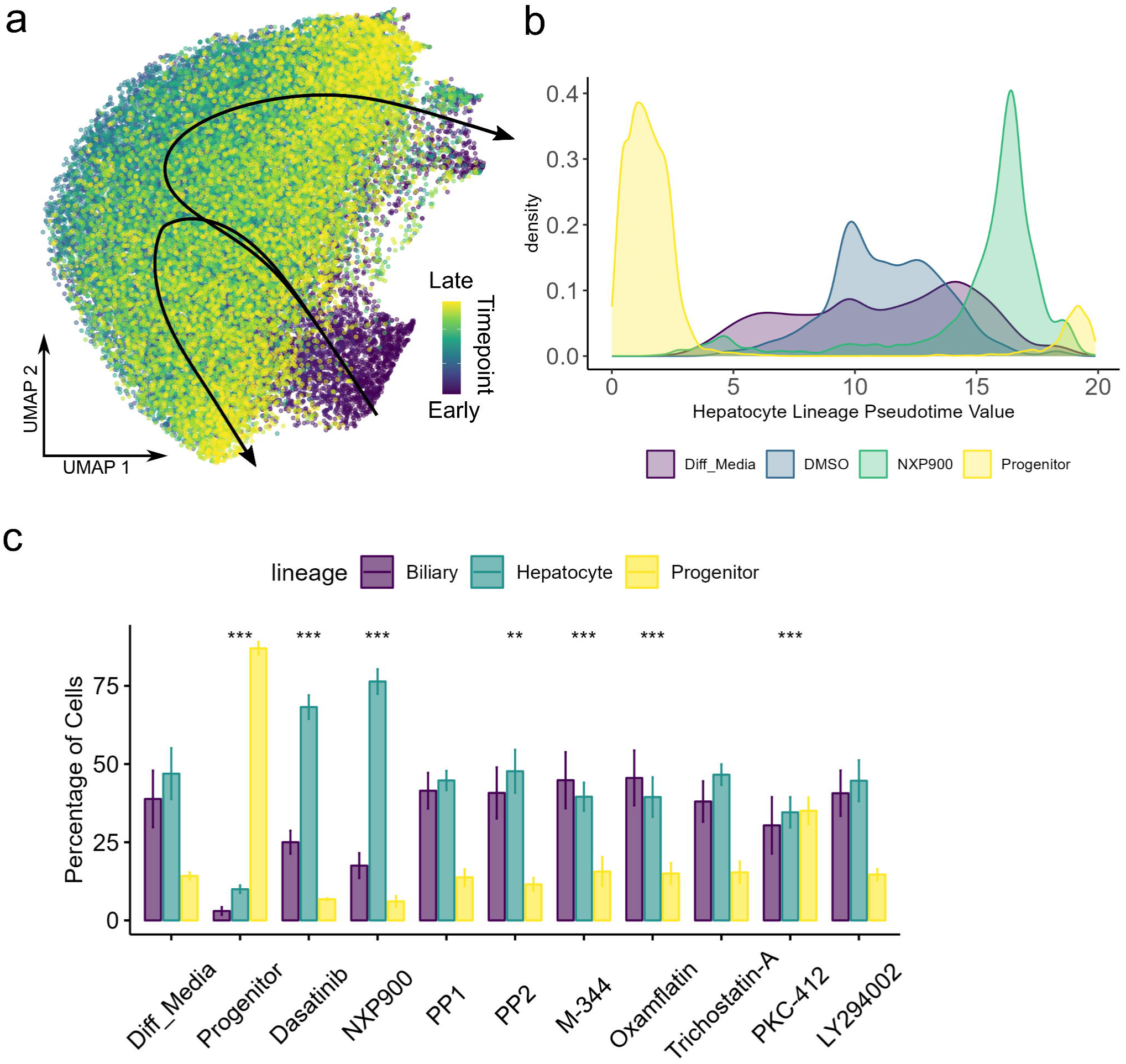
Single-cell morphological trajectory analysis. a) Slingshot trajectory projected onto UMAP embedding of HepaRG^®^ differentiation atlas. b) Density plot showing cell distributions along the hepatocyte lineage trajectory for controls and NXP900 treatment. c) Percentage of cells belonging to each lineage across compound treatments. (Mean ±SD). Chi square test, reference distribution= differentiated control, ** p<0.01, *** p<0.001, n =3.

Strikingly in the image level analysis, the top concentration of NXP900 (10 uM) appears to deviate from the desired phenotype since the Euclidean distance to the positive control increases (**Fig. 2d**). In multiparametric high-content phenotypic profiling studies such deviations at high concentrations are common and are often thought to be a consequence of cytotoxicity. However, the single-cell analysis reveals that this is because NXP900 treatment has dramatically altered the cell type composition, highly enriching the hepatocyte-like population (**Fig. 4c**). This highlights the additional information and resolution achievable when using the single-cell level data in comparison to the industry standard image level analysis of drug treatments. Further the SFK inhibitor NXP900, and Dasatinib to a certain extent, appear to promote hepatic differentiation in the 8 day assay to the same extent as the final timepoints in the timecourse data when the cells have had the full 4 weeks to differentiate, therefore outperforming the differentiated controls in the assay, indicating that the SFK inhibitors significantly accelerate the hepatocyte differentiation process.

### Trajectory Inference

In order to quantify biological progression through this process of bilineage differentiation, we computationally inferred the differentiation trajectory of liver cells from the single-cell Cell Painting atlas by pseudo-time mapping using the slingshot method ^30^. This produced two lineage trajectories starting from the single-cell progenitor population as expected (**Fig. 5a**). We then projected the NXP900 single-cell data onto the liver differentiation trajectories to determine the pseudotime scores and quantitatively compare NXP900 induced differentiation with the controls. NXP900 led to much greater pseudotime values for the hepatocyte lineage when compared to both the undifferentiated control and the differentiated control (commercially available supplement, see methods), supporting the idea that NXP900 is promoting the hepatocyte-like cells in the HepaRG^®^ line, and further, promoting a more mature hepatocyte-like cell than the differentiated control (**Fig. 5b**). To quantitatively compare how the compounds affected the ratios of the two mature cell types we used the lineage weightings to assign each cell to either the progenitor (pre-branching of the two lineages), the hepatocyte, or the biliary lineage. From this we can see that the majority of the progenitor control cells map to the portion of the trajectory that is pre-branching (progenitors), while the differentiated control and most of the hit compounds lead to a nearly 50:50 split of the biliary and hepatocyte lineages (**Fig. 5c**). NXP900 and Dasatinib cause a significant increase in cells along the hepatic lineage, increasing the proportion to over 75% in the NXP900 treated populations. This could have important implications in minimising heterogeneity *in vitro* and enabling more efficient engraftment *in vivo*.

### Validation of hits in primary human hepatic progenitor cells

Our morphological single-cell trajectory inference of small molecule compound perturbations suggests that SFK inhibitors promote the hepatocyte-like lineage in the HepaRG^®^ cells, something that cannot be detected using the image level data. Small molecules that can be added to culture media as a supplement to promote the functional maturity of primary hepatocytes *in vitro* are of great interest to the liver regeneration community. We therefore chose to test the SFK inhibitors in primary adult human hepatic progenitor cells (HPC), bipotent cells isolated from discarded donor organs, to measure multiple functional hepatic markers and confirm the ability of the compounds to induce differentiation of human HPC towards a hepatocyte-like phenotype. The following markers, HNF4a (an essential transcription factor involved in the functional differentiation of hepatocytes during development), CYP2E1 (cytochrome P450 enzyme) and albumin (Essential role in transportation) were selected to infer hepatocyte differentiation. Biliary epithelial cell marker, CK7 and proliferation marker Ki67 were also included in the phenotypic panel. Incubation of human HPC with the SFK inhibitors NXP900, PP1 and PP2 all demonstrate a significant increase in the proportion of cells expressing Cyp2E1 and HNF4a compared to untreated controls. Albumin expression remained low across all compound treatments, suggesting only partial differentiation at this timepoint. Of note, PP1 also significantly reduced the biliary epithelial marker CK7, suggestive of a phenotypic shift (**Fig. 6a and b**).

**Fig 6.**
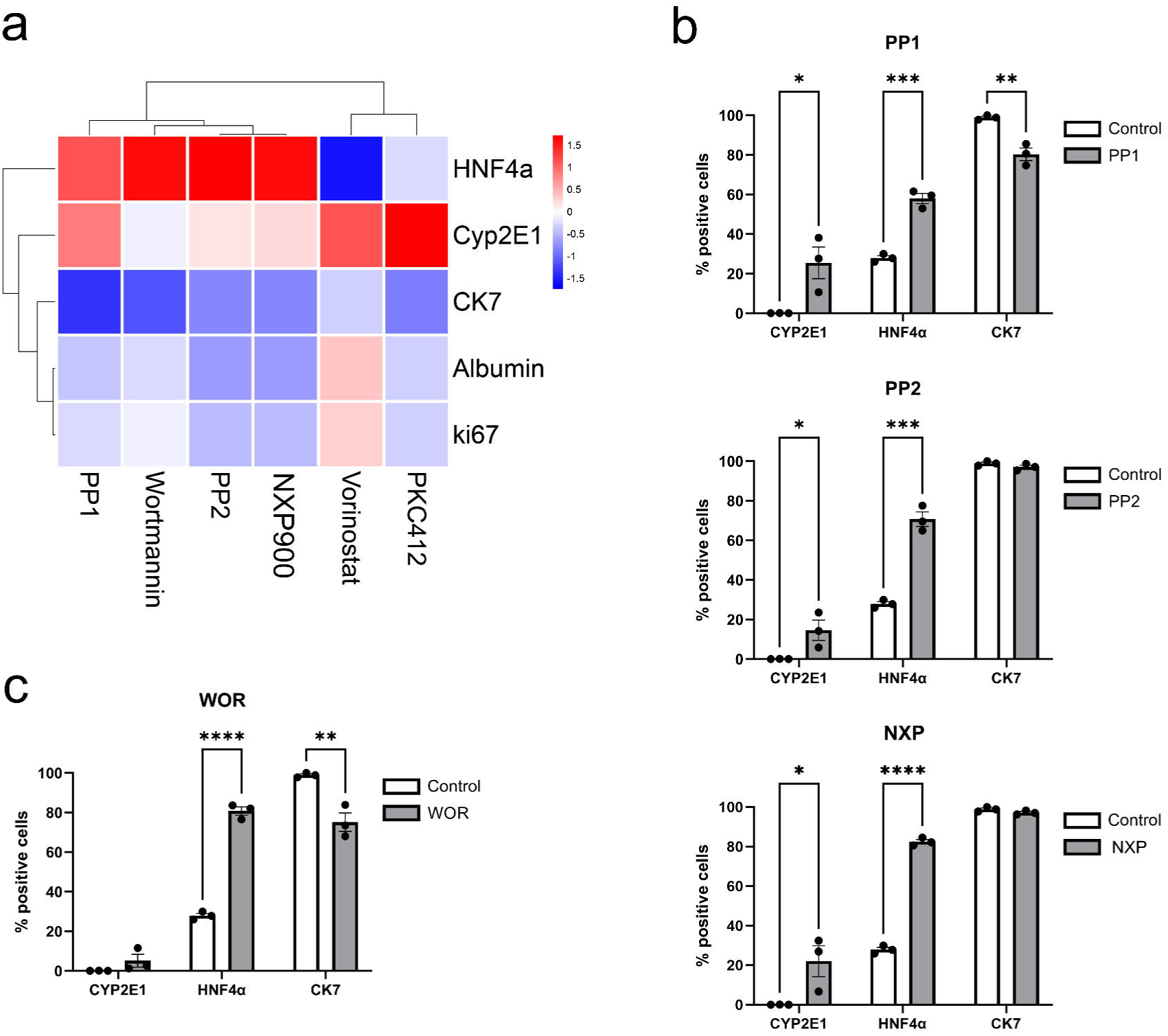
Validation of candidate hepatocyte-like inducing small molecules in primary human HPC. A) Heatmap of phenotypic marker expression in response to compound treatments. Data normalized to control expression. B) Percentage of cells positive for hepatocyte markers CYP2E1 and HNF4a and biliary epithelial marker CK7 in response to SFK inhibitors. (Mean ±SEM) Multiple t-tests, * p < 0.05, ** p < 0.01, *** p < 0.001, n = 3 per group. C) Percentage of cells positive for hepatocyte markers CYP2E1 and HNF4a and biliary epithelial marker CK7 in response to PI3K-AKT inhibitor, Wortmannin. (Mean ±SEM) Multiple t-tests, ** p < 0.01, **** p < 0.001, n = 3 per group.

We further tested several other hits from the HepaRG^®^ screen in the primary human HPC and found that several hits increased a single hepatocyte functional marker (**Fig. 6a**). Interestingly, the PI3K-AKT inhibitor, Wortmannin demonstrated a 40% increase in HNF4a expression coupled with a reduction in CK7 expression (**Fig. 6c**), suggestive of differentiation.

Taken together, these data validate the use of single-cell morphological trajectory inference assay to identify candidate small-molecule modulators of liver cell differentiation for further testing in primary human cells. Of particular interest would be the use of SFK inhibitor NXP900 as well as the PI3K-AKT inhibitor, Wortmannin.

## Discussion

During development, in response to endogenous and exogenous stimuli, disease onset, and throughout life, cells transition from one functional “state” to another ^31^. Understanding and controlling cell state transitions and long-term cell fate has important applications in therapeutic development for various developmental disorders and disease areas. There are many important markers of a cells physiological state including gene expression and cellular morphology ^32^ providing opportunities for molecular and phenotypic profiling technologies to track cell state transition and long-term fate decisions in biological samples.

Single-cell genomic technologies such as single-cell RNA-Sequencing (scRNA-Seq) ^33^ has significantly advanced in-depth mechanistic studies of cell fate and dynamic cell state transitions in biological samples ^34,35^. Recent advances in scRNA-Seq and computational trajectory inference methods have been applied to infer the dynamics of cell fate across a variety of complex biological models and samples ^30,36–39^. Such studies include applications of scRNA-Seq to spatially map gene expression and cell phenotypes across the entire mammalian liver ^40^ and understand the progression of liver diseases such as cirrhosis ^41^ and non-alcoholic steatohepatitis (NASH) ^42^. High-throughput application of scRNA-seq technology, however, is restricted by costs limiting its use in early-stage drug discovery for target identification and *in vitro* or *in vivo* proof-of-concept studies on promising leads or drug candidates. In contrast, multiparametric high-content imaging technology is more cost effective per sample and can generate morphological phenotypic fingerprints at single-cell level at scale ^43^. Thus, in addition to target discovery and proof-of-concept studies, high-content morphological profiling is also applicable to systematic phenotypic screening of large sets of target-annotated and diverse chemical libraries and/or druggable whole genome targeting siRNA and CRISPR libraries to enable a comprehensive survey of chemical and biological modulators of cell states.

The morphological trajectory inference described here represents a holistic, high-throughput and inexpensive alternative to antibody based functional hepatocyte assays and RNA-sequencing, providing deep insights into the impact of small-molecule perturbations on liver cellular differentiation. Further, by enabling the computational alignment and modelling of cellular states this single-cell Cell Painting analysis pipeline has broad reaching applications offering a new way to quantify single-cell responses across perturbation screens, disease progression, and cellular differentiation to name a few. This technique will be particularly powerful when applied to multi-cellular assays as exemplified here in the mixed lineage analysis where cell non-autonomous or cell-type specific responses to perturbations can be assessed within the context of complex multi-cell assays. Further since the analysis strategy models biology as a continuous trajectory, we can triage for off-target and unwanted trajectories caused by compounds while previously we have only been able to assess perturbations in terms of distance to static phenotypes of interest. This high-throughput compatible assay therefore has an incredible resolution for not only identifying cell type specific and biologically relevant hits but we can also begin to unravel cell state transitions, cell dynamics, and lineage specificity.

In this work we have identified a group of SFK inhibitors that promote differentiation of liver progenitor cells towards a hepatocyte-like phenotype *in vitro* across two liver cell models, the HepaRG^®^ liver progenitor cells and primary human hepatic progenitor cells. NXP900 increases the hepatic-like population of HepaRG^®^ cells to over 75%. This enrichment for hepatic cells is a very useful property since it will minimise heterogeneity and subsequent co-culture effects impacting on *in vitro* model performance. It could also potentially reduce asynchrony in the differentiation process which is an important consideration *in vitro* and *in vivo*. Further, this could be important in the context of cell transplant for efficient cell engraftment and tissue integration to rapidly boost hepato-cellular function and improve host liver performance.

SFK activity and downstream signalling has previously been implicated in the progression of a number of liver disorders including hepatocellular carcinoma ^44^ and liver fibrosis ^45^. In addition, a role for c-SRC in cell fate determination during endodermal commitment of human iPSCs has recently been described ^46^, the kinase inhibitor PP2 has previously been shown to protect against Keratin mutation-induced mouse liver injury through Src inhibition (Li et al., 2023), and previous studies have demonstrated that genetic deletion of Src’s binding partner focal adhesion kinase (FAK) accelerates liver regeneration ^48^. It is our understanding that ours is the first study to demonstrate a role for small molecule SFK inhibitors in the promotion of hepatocyte progenitor differentiation and as a potential therapeutic opportunity for liver regeneration.

In contrast to currently approved dual Src-Abl inhibitors, including dasatinib, which block SRC in the active “open” conformation (promoting the association of Src and signalling partners via allosteric facilitation) ^49^, NXP900 binds and locks Src tyrosine kinase in a closed inactive conformation, inhibiting both scaffolding and catalytic activity ^50^. The novel mechanism of action of NXP900 contributes to its unique target selectivity profile with 1000-fold selectivity for SFK members over ABL kinase and results in highly potent, selective and sustained pathway inhibition, *in vitro* and *in vivo* ^50^. Dasatinib is already FDA approved and NXP900 has progressed into phase 1 clinical studies and thus each represent promising drug candidates to stimulate endogenous or exogenously administered (i.e., cell therapy) liver cell differentiation in situ to restore damaged liver function. The SFK inhibitors could also be used as a cell culture additive to maintain liver differentiation *in vitro* for biological investigation and drug discovery applications.

In summary, we describe a novel application of trajectory inference to high-throughput morphological data to quantify cell state transitions at the single-cell level in a high-throughput screening format. We have demonstrated the utility of the method by application to a small molecule compound library screen in a liver progenitor differentiation assay and validated the hits by confirming activity in lower throughput primary human hepatic progenitor differentiation assays. This demonstrates our assay is validated in a high-throughput format and suitable for screening larger libraries of target-annotated compounds and diverse chemicals to identify additional therapeutic targets and chemical starting points and thus launch new drug discovery programs for liver regeneration. The validated hits from the pilot screen include a series of SFK inhibitors revealing a novel role for SRC (and other SFK members) in accelerating hepatocyte differentiation. Future work to demonstrate the clinical utility of SFK inhibitors in promoting liver regeneration includes *in vitro* functional assays such as drug metabolism and albumin secretion. Further, *in vivo* studies to investigate the effects on efficiency of expansion and differentiation of exogenously applied hepatocytes as well as the ability to directly target endogenous HPC repair mechanisms will be important. Finally, single-cell morphological trajectory inference is applicable to other high-content image-based assays, particularly stem cell differentiation and disease progression and thus has broad utility for comprehensive investigation of modulators of cell state transitions.

## Methods

### HepaRG cell culture

Cells were supplied by Biopredic International and cultured according to the user guide. Briefly, cells were thawed and subcultured to create Master and Working banks of frozen cells. For this cells were passaged at day 7 post thaw and then after that cells were passaged every 14 days for expansion and frozen down. Cells were cultured in William’s E medium supplemented with GlutaMAX (2 mM) and HepaRG^®^ Growth Medium Supplement (ADD711C, Biopredic International). The medium was changed every two to three days.

### Screening

Compound libraries: A collection of 496 known bioactive compounds included the ENZO SCREEN-WELL® kinase (80 compounds), protease (53 compounds) and epigenetic (43 compounds) libraries and a subset of FDA/EMA approved drugs representing broad pharmacological diversity of all approved small molecules (320 compounds) (**Supplementary Table 1**). Compound handling: Compound source plates solubilized in DMSO were made at 1,000-fold assay concentration in 384-well plates and transferred to the cells using a Biomek FX liquid handler with an overall dilution in media of 1:1000 from source to assay plate (final DMSO concentration =0.1%DMSO).

The primary screen was carried out as a single replicate at two compound concentrations (5uM and 0.5uM) and the validation dose response study was in triplicate.

For screening, HepaRG^®^ cells were used between passages 16 and 18. Cells were thawed and seeded in T175 flasks at 5.5 million cells per flask and incubated for a week with media changes every 2-3 days. After a week the cells were trypsinised and seeded at confluency, 20,000 cells per well, in 384-well plates (#781091, Greiner). After 72 hrs the media was changed (70% media change) and compounds and positive control differentiation supplements (ADD721C, Biopredic International) were added. Drug incubations were for 96hrs in total with a media change and drug addition after 48 hrs. 24 hrs before the end of the assay progenitor control HepaRG^®^ cells were spiked into an empty column at 3000 cells per well to allow normalisation to the timecourse dataset. Plates were fixed in 4% formaldehyde, stained (see Cell Painting section below) and imaged (see Image Acquisition section below).

### Timecourse

For the untreated, time-course work, cells were seeded at 3000 cells per well into 20x 384-well plates (#781091, Greiner), 5 columns per plate. Media was changed (70%) every two to three days. One plate was fixed every 24-72hrs in 4% formaldehyde for up to 28 days following cell seeding (**Table 1**). For the plates remaining after two weeks, the supplement was changed to the HepaRG^®^ Differentiation Medium Supplement without antibiotics (ADD721C, Biopredic International) and media changes continued until all plates had been fixed.

**Table 1.**
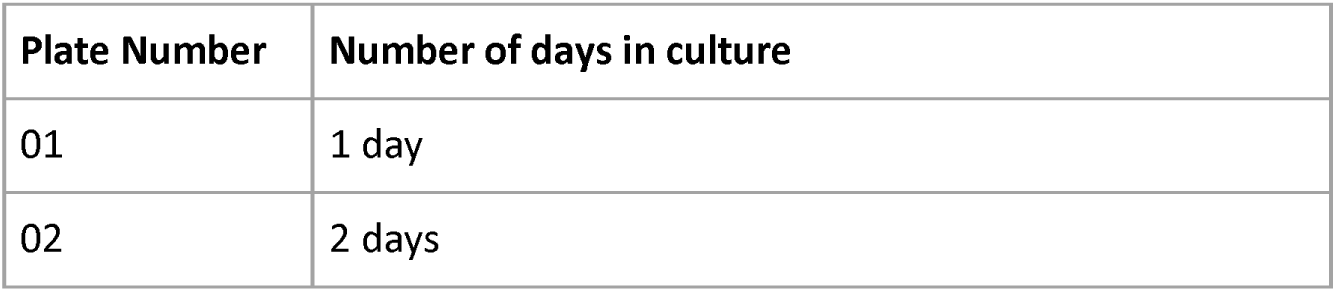
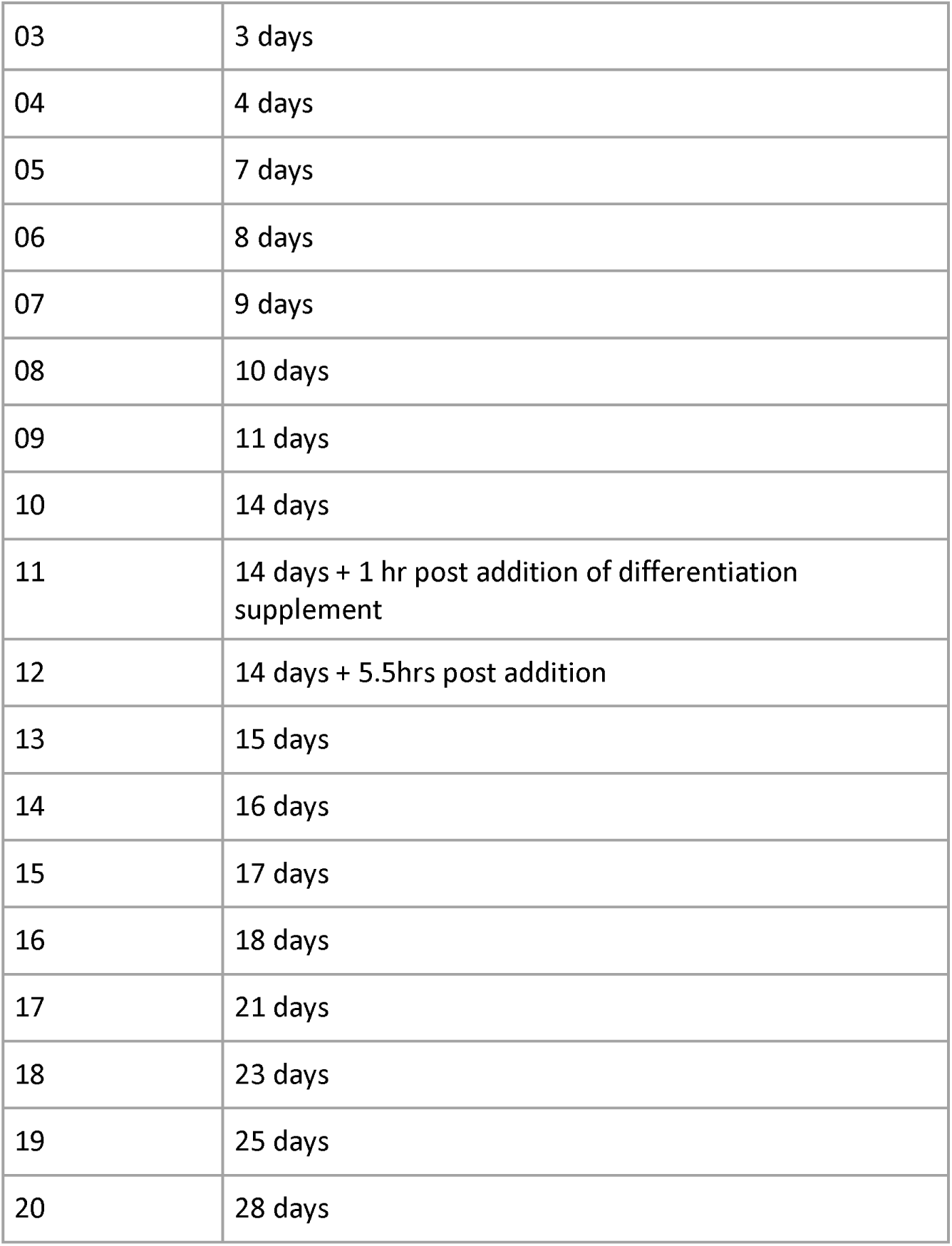
Plate number and days in culture after seeding.

| Plate Number | Number of days in culture |
| --- | --- |
| 01 | 1 day |
| 02 | 2 days |
| 03 | 3 days |
| 04 | 4 days |
| 05 | 7 days |
| 06 | 8 days |
| 07 | 9 days |
| 08 | 10 days |
| 09 | 11 days |
| 10 | 14 days |
| 11 | 14 days + 1 hr post addition of differentiation supplement |
| 12 | 14 days + 5.5hrs post addition |
| 13 | 15 days |
| 14 | 16 days |
| 15 | 17 days |
| 16 | 18 days |
| 17 | 21 days |
| 18 | 23 days |
| 19 | 25 days |
| 20 | 28 days |

### Cell Painting

Cell Painting was carried out according to the published protocol ^15^with the following exceptions: Mitotracker DeepRed was added post fixation in all assays (screen, validation and timecourse), to allow a single staining solution to be added to all timecourse plates after fixation of the final plate to avoid batch effects in the timecourse data.

After fixation, plates were washed twice with PBS and the staining solution (**Table 2**) was added containing 0.1% Triton-X100 in 1% bovine serum albumin (BSA). The Syto14 stain was omitted in case a cell type specific marker was required in future. The staining solution was added to each well (25 µL) and incubated in the dark at room temperature (30 minutes), followed by three washes with PBS and no final aspiration. Plates were foil sealed.

**Table 2.** Cell Painting reagent information.

| Stain | Structure | Wavelength<br>(ex/em [nm]) | Channel | Assay<br>Concentration | Cat No;<br>Supplier |
| --- | --- | --- | --- | --- | --- |
| Hoescht 33342 | Nuclei | 387/447 | DAPI | 6 µg/mL | #H1399; Mol.<br>Probes |
| Phalloidin 594 | F-actin | 562/624 | TxRED | 0.14X | #ab176757;<br>Abcam |
| Wheat germ<br>agglutinin Alexa<br>Fluor 594 | Golgi and Plasma<br>Membrane | 562/624 | TxRED | 1.5 µg/mL | #W11262;<br>Invitrogen |
| Concanavalin A<br>Alexa Fluor 488 | Endoplasmic<br>Reticulum | 462/520 | FITC | 40 µg/mL | #C11252;<br>Invitrogen |
| MitoTracker<br>DeepRed | Mitochondria | 628/692 | CY5 | 600 nM | #M22426;<br>Invitrogen |

### Image Acquisition

Plates were imaged on an ImageXpress Confocal (Molecular Devices, USA) equipped with a robotic plate loader (Scara4, PAA, UK). Four fields of view were captured per well using a 20x objective and four filters (**Table 2**).

### Image Analysis

CellProfiler v4.2.1 software was used to segment the cells and extract 1004 features per cell per image. First the pipeline identified the nuclei from the DAPI channel and used these as seeds to aid a segmentation algorithm to identify the cell boundaries from the TxRed channel, and finally these two masks were subtracted to give the cytoplasm. These three masks marking the cellular boundaries were then used to measure morphological features including size, shape, texture, and intensity per object across the five image channels.

The median and standard deviation values were calculated for all features per image, leading to 2008 features total. StratomineR software was used for feature selection, normalisation, feature scaling, dimension reduction, distance, and similarity metrics. 848 features were used in the analysis. Plates were normalised to the samples on them and then features scaled using robust z-scores. Principal component analysis was applied for dimension reduction and hits were defined based on Euclidean and Pearson distances.

A Random Forest classifier was implemented on the three classes of controls (Progenitors, Undifferentiated, Differentiated) using R’s RandomForest package with the default parameters. There were 64 data points in each class. 28 variables were used at each split and 500 trees were used.

Validation analysis was similar except only median feature values were used and after feature selection 189 features were used in the analysis. Plates were normalised to the negative controls.

### Single-cell analysis

The single-cell timecourse and drugs of interest datasets were initially filtered based on known cellular size parameters -nuclear area of greater than 500 and less than 3250 px^2^ and a total cell area of less than 37000 px^2^. These two datasets were down sampled to 15000 cells per timepoint/treatment and standardised separately using the progenitor controls before combining into one dataset (788 morphological features). Principal component analysis followed by UMAP on the top 10 components were used for visualisation of the combined single-cell data. Progenitor cells (24 hrs post seeding) overlay perfectly across the two datasets demonstrating no need for single-cell batch correction (**Supplementary Fig. 4a**).

Doublet cells (missegmentation of two cells as one cell) formed a distinct cluster in the data and were removed by training a balanced 3 class random forest classifier (Progenitor (100 annotated cells), Doublet (100 annotated cells), Inlier (100 cells from the bulk of liver cells between progenitors and fully differentiated) (**Supplementary Fig. 4b**) to label and remove doublets.

Raw features (788) were restandardised for the cleaned datasets as before and principal component analysis and UMAP were repeated for visualisation. An extreme outlier cluster of 66 artefacts was removed and the standardisation, principal component analysis and UMAP were carried out on this final clean dataset.

### Trajectory inference

We used the R package Slingshot ^30^ to perform single-cell trajectory analysis. The single-cell UMAP dataset generated above was separated into the timecourse data and the drug treated data. The timecourse data was clustered using Gaussian mixture modelling implemented in the r package mclust ^51^. Using Slingshot to generate the trajectories we first defined the backbone by selecting the clusters that represented the progenitor starting cells and the two differentiated cell types. Slingshot was passed the first two UMAP coordinates, the single-cell cluster labels, and the trajectory backbone clusters (start and end points).

The drug treated single-cell data was then projected onto the timecourse trajectories. Specifically, the UMAP embedding of the drug data was used to determine the positions of the cells along the previously constructed trajectories using the predict function in slingshot, producing a hybrid object with the trajectories from the timecourse data, but the pseudotime values and weights for the drug treated cells.

Multiple runs of the single-cell trajectory analysis produced similar embedding, clustering and trajectory results.

Single-cell lineage assignment was generated from the slingshot lineage weightings. Cells were assigned “Hepatocyte” if they had a lineage 1 weighting equal to 1, a lineage 2 weighting of less than 1, and a pseudotime value of greater than 3 (branching timepoint), “Cholangiocyte” labels were assigned for the opposite lineage weightings, and the “Progenitor” label was assigned to the remaining cells (pseudotime value 3 or less).

### Primary human HPC differentiation assay

#### Cell culture and compound treatment

Primary human hepatic progenitor cells were isolated from discarded donor organs as previously described (Hallett et al. 2020). Briefly, the donor liver is chopped into small pieces before undergoing enzymatic digestion and CD133 selection via magnetic activated cell sorting (Miltenyi Biotec, Cat#170-076-719). Primary human HPC are routinely expanded as 3D organoids suspended in Matrigel (Corning) and cultured in expansion media; AdDMEM/F12 (Invitrogen) base media, supplemented with Pen-Strep (Invitrogen, 15140-122), HEPES buffer (10mM, Gibco) Glutamax (Life Technologies), B27(Life Technologies), N2 (Life Technologies), N-acetylcysteine (1.25mM, Sigma-Aldrich), Nicotinamide (10mM, Sigma-Aldrich), gastrin (10nM, Sigma-Aldrich), A83-01 (5μM, Miltenyi Biotec), the growth factors: FGF10 (100ng/mL, Perprotech), HGF (25ng/mL, R&D Systems), EGF (50 ng/ml, Peprotech), Rspo1 (500ng/mL, Peprotech), Forskolin (10 μM) and the ROCK inhibitor: Y-27632 (10μM, Miltenyi Biotech).

Prior to compound screening, 3D organoids were dissociated into single-cells and seeded on laminin-521 coated plates to expand as 2D cells. Organoids are washed in cold PBS-EDTA (PE; 10mM, Gibco) and broken down with 1U/mL Dispase for 30 minutes at 37°C. Organoids are further disrupted via manual pipetting and incubated for 30 minutes. Once broken down, the cells are washed with cold PE and incubated with a TrypLE mice solution (10X TrypLE + 1X TrypLE at 1:4) for 5 minutes with gentle agitation at 37°C to obtain single cells. Cells were plated on laminin-521 (10 µg/mL) coated plates, cultured in expansion media, replenished every 3 – 4 days.

To screen candidate compounds, 2D human HPC were seeded on to 96-well plates (2 x 10^4^ cells/well) and allowed to attach for 24 hours. The compounds and final concentrations selected for validation in human HPC are outlined in **Supplementary Table 3**. The selected compounds were administered along with fresh expansion media every 48 hours. After 7 days, cells were immunostained for a selection of phenotypic markers and counterstained with Cell Mask Deep Red (cell membrane) and DAPI (nuclei) (**Table 3**).

**Table 3.** Primary human HPC phenotypic panel information.

| Stain | Phenotypic marker | Cell fixation method | Assay Concentration | Cat No; Supplier |
| --- | --- | --- | --- | --- |
| CK7 | Biliary epithelial cell (HPC) | 1:1 methanol acetone | 0.36 µg/mL | #ab68459; Abcam |
| HNF4α | Hepatocyte | 10% formalin | 3.3 µg/mL | #PP-H1415-OC; R&D Systems |
| Albumin | Hepatocyte | 1:1 methanol acetone | 3.3 µg/mL | #ab2406, Abcam |
| CYP2E1 | Hepatocyte | 1:1 methanol acetone | 0.3 µg/mL | #HPA009128, Atlas Antibodies |
| Ki67 | Proliferation | 10% formalin | 0.1 µg/mL | #ab1667, Abcam |
| Cell Mask | Cell membrane | n/a | 0.5 µg/mL | #C10046,<br>ThermoFisher |
| DAPI | nuclei | n/a | 1 µg/mL | #D9542,<br>Scientific<br>Laboratory<br>Supplies |

#### Immunocytochemistry

Prior to immunostaining, human HPC were fixed with 1:1 methanol acetone or 10% formalin for 20 minutes. Cells underwent permeabilization with 0.1% Triton X-100 (Sigma) in PBS for 15 minutes, followed by protein block (Abcam) for 30 minutes and were then incubated with relevant primary antibodies diluted in antibody diluent (Abcam) at 4°C overnight. Cells were finally stained with Alexa Fluor 488 secondary antibodies (Invitrogen) combined with Cell Mask and DAPI for 30 minutes at room temperature.

#### Image acquisition

Images were captured by the Opera Phenix High Content Screen System and analyzed by Harmony software. Total cell numbers were determined by combined detection of cell membrane and nuclei. Cells positive for individual phenotypic markers were identified utilizing expression thresholds over relevant isotype controls and expressed as a % of positive cells per well (performed in triplicate).

## Data Availability

Data will be made available upon manuscript publication. Code will be made available on our Github page upon manuscript publication.

## Acknowledgments

This work was supported by the UK Medical Research Council - UK Regenerative Medicine Platform. We thank Justyna Cholewa-Waclaw and the High Content and Imaging core facility at Institute for Regeneration Repair for their technical help with experimental design, data acquisition and analysis. REG was funded by MRC Transition Fellowship.

## Author Contributions

R.E.G contributed to the conceptualisation, and acquisition, analysis, and interpretation of the data, as well as drafting the manuscript. R.Z contributed acquisition and analysis of the data. J.W contributed designing the analysis. A.U.B provided reagents and contributed to interpretation of the data and manuscript revision. D.C.H contributed conceptualisation of the work. S.J.F contributed conceptualisation of the work as well as interpretation of the data. V.G contributed design of the work and acquisition, analysis, and interpretation of the data, as well as drafting the manuscript. N.O.C contributed conceptualisation and drafting the manuscript. All authors contributed to the discussion of the results and approved the final version of the manuscript.

## Competing interests

A.U-B and N.O.C disclose the following patents, EP3298015B1, JP6684831B2, US10294227B2, CN107849050B, and CA3021550A1 pertaining to the discovery of eCF506/NXP900 that have been licensed to Nuvectis Pharma Inc. A.U-B and N.O.C hold grants from Nuvectis Pharma to study eCF06/NXP900 outside of the submitted work. D.C.H is a co-founder, director and shareholder in Stimuliver ApS and Stemnovate Limited.

## Materials and Correspondence

Correspondence to Rebecca E Graham and Neil O Carragher.

