## Supplementary Data for "Single-cell morphological tracking of liver cell states to identify small-molecule modulators of liver differentiation"

### Slide 1
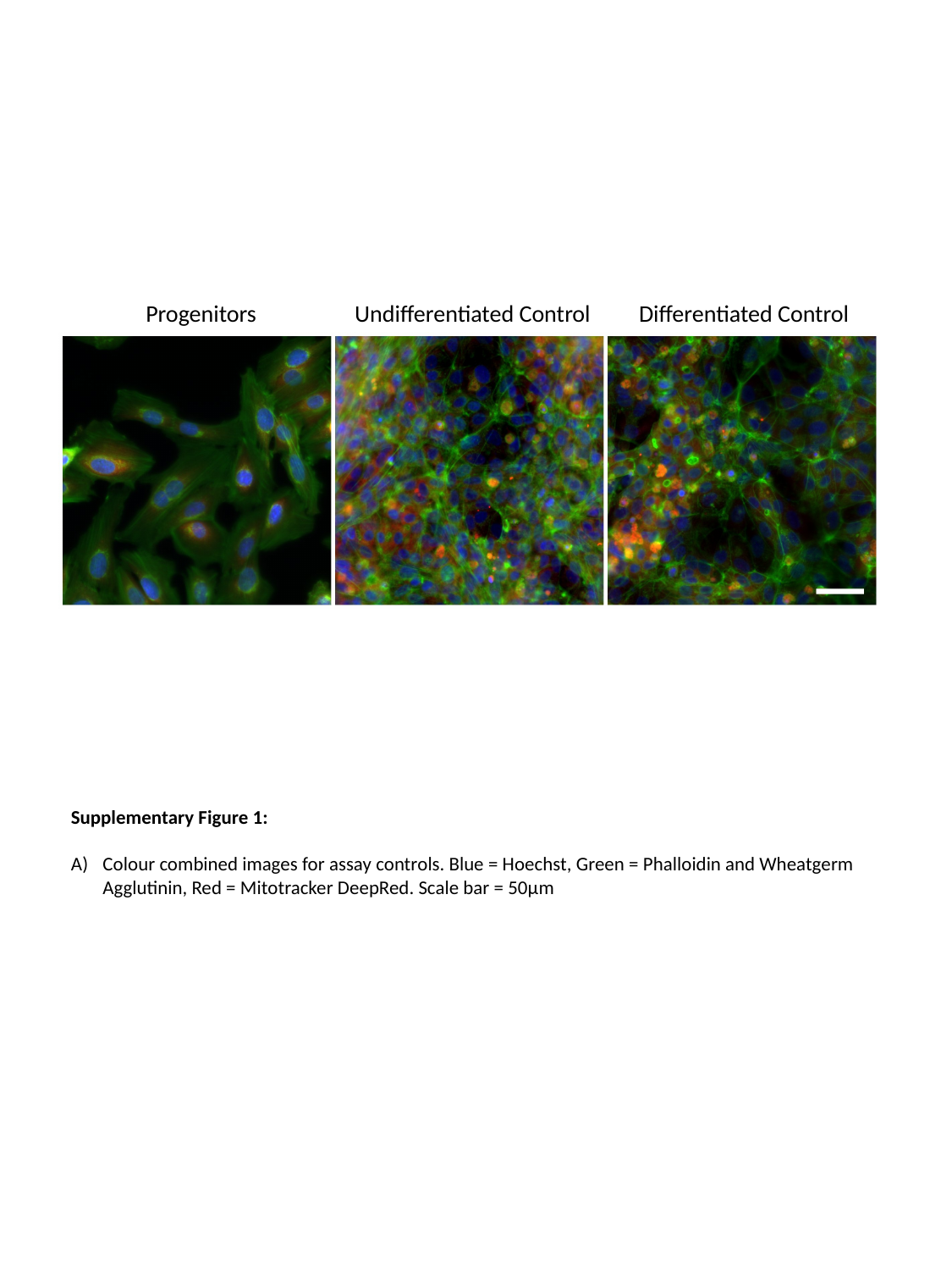

| Progenitors | Undifferentiated Control | Differentiated Control |
| --- | --- | --- |
Supplementary Figure 1:
Colour combined images for assay controls. Blue = Hoechst, Green = Phalloidin and Wheatgerm Agglutinin, Red = Mitotracker DeepRed. Scale bar = 50µm

### Slide 2
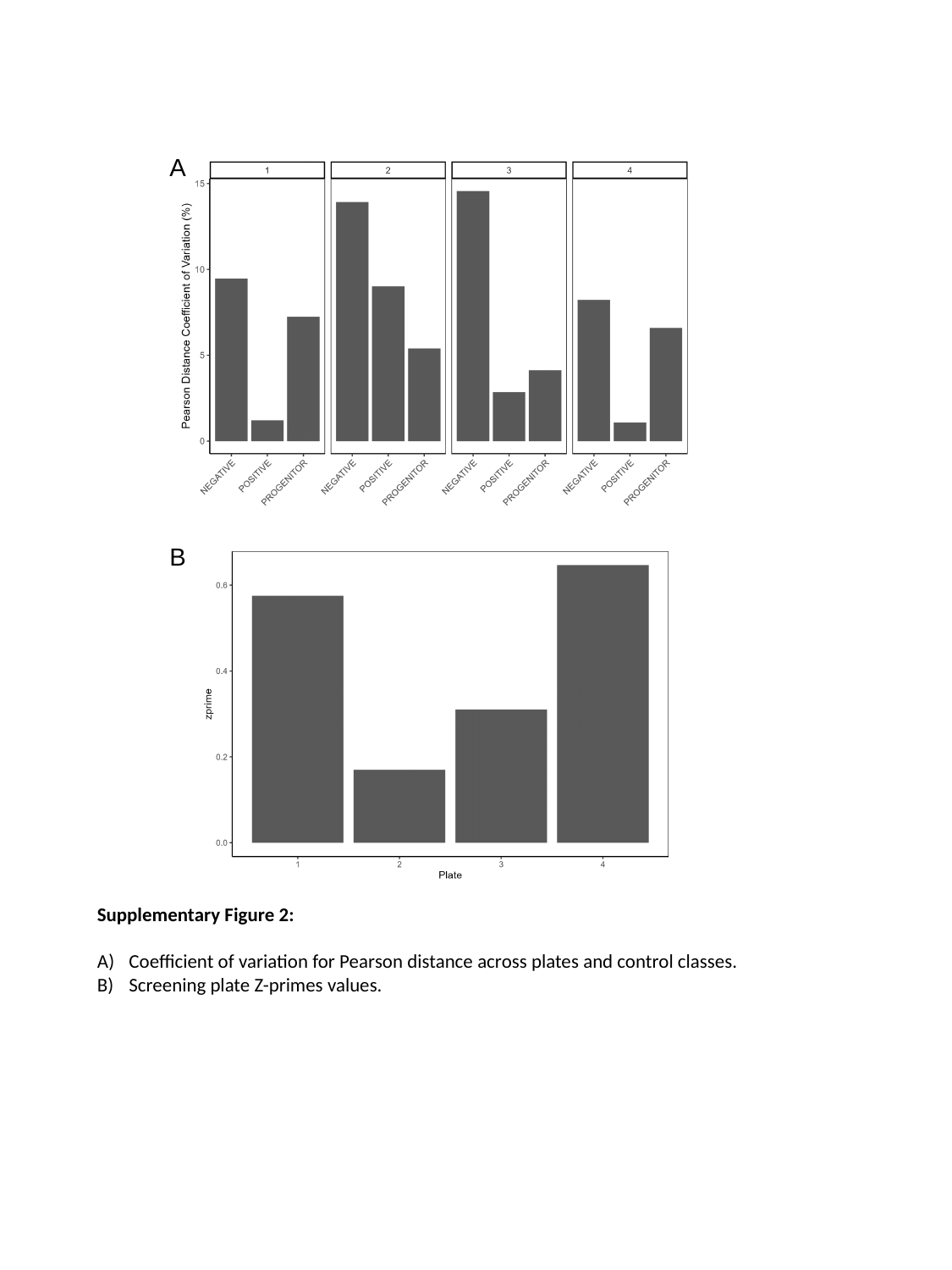

A
B
Supplementary Figure 2:
Coefficient of variation for Pearson distance across plates and control classes.
Screening plate Z-primes values.

### Slide 3
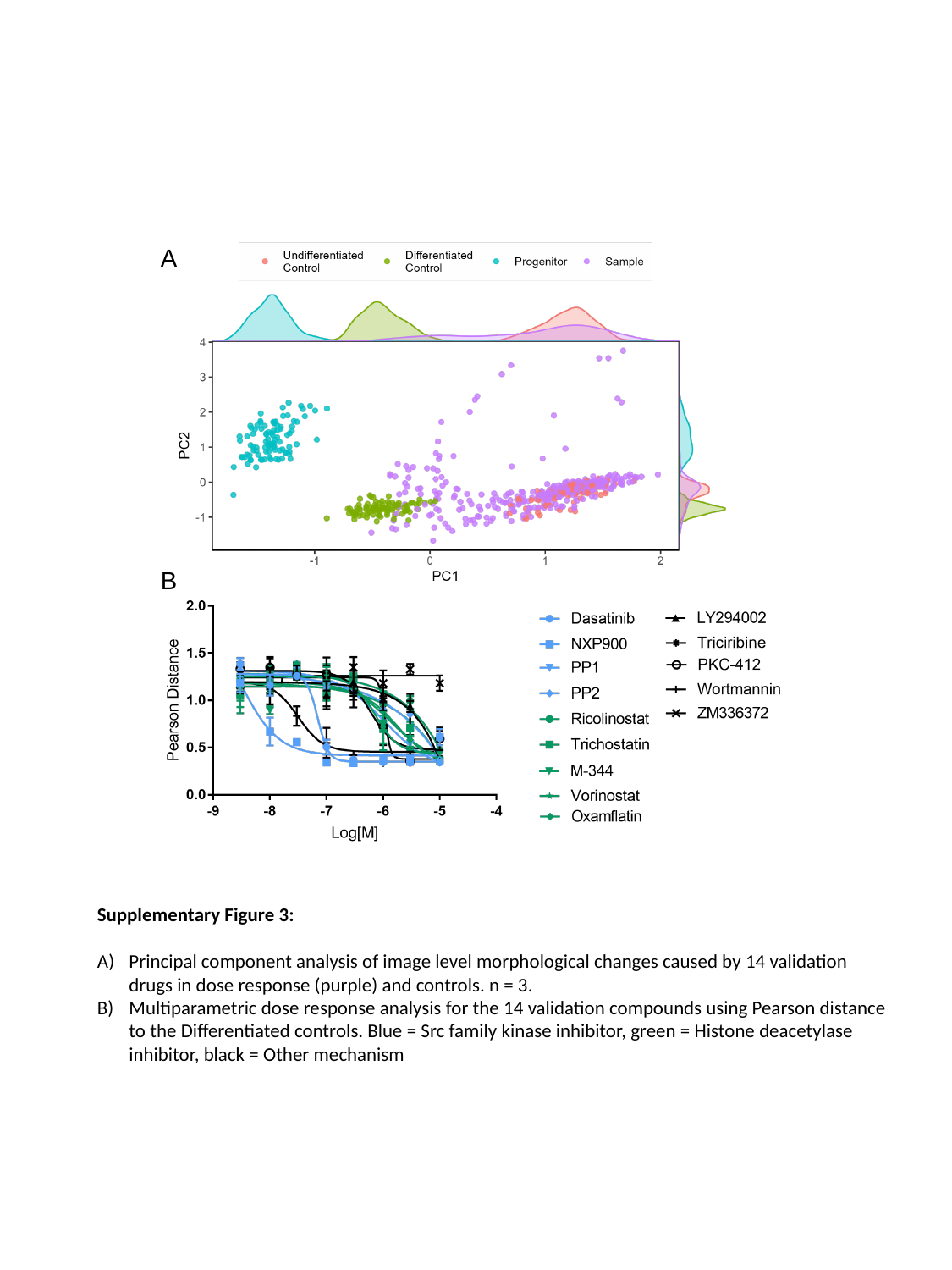

A
B
Supplementary Figure 3:
Principal component analysis of image level morphological changes caused by 14 validation drugs in dose response (purple) and controls. n = 3.
Multiparametric dose response analysis for the 14 validation compounds using Pearson distance to the Differentiated controls. Blue = Src family kinase inhibitor, green = Histone deacetylase inhibitor, black = Other mechanism

### Slide 4
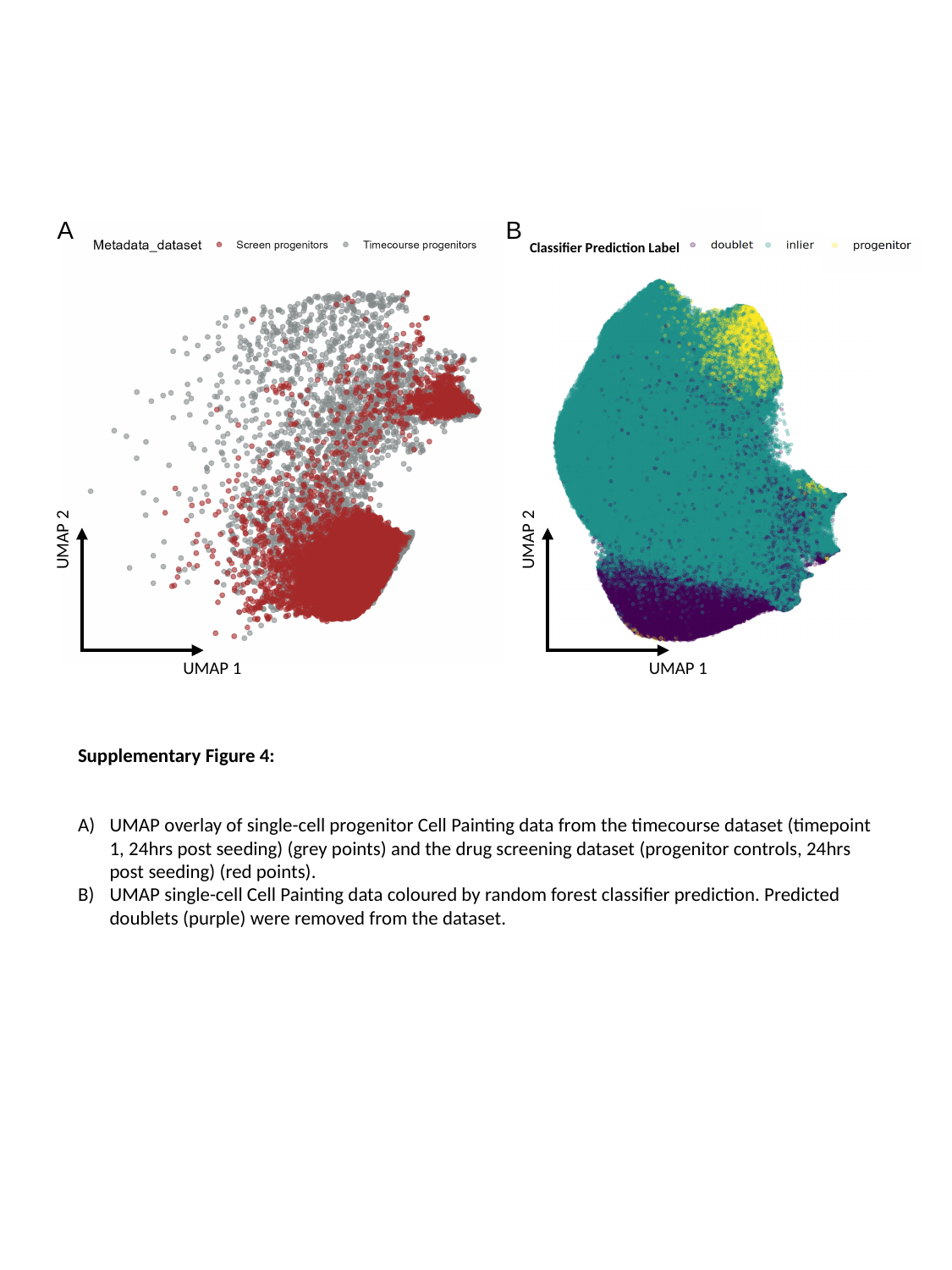

A
B
Classifier Prediction Label
UMAP 2
UMAP 2
UMAP 1
UMAP 1
Supplementary Figure 4:
UMAP overlay of single-cell progenitor Cell Painting data from the timecourse dataset (timepoint 1, 24hrs post seeding) (grey points) and the drug screening dataset (progenitor controls, 24hrs post seeding) (red points).
UMAP single-cell Cell Painting data coloured by random forest classifier prediction. Predicted doublets (purple) were removed from the dataset.

### Slide 5
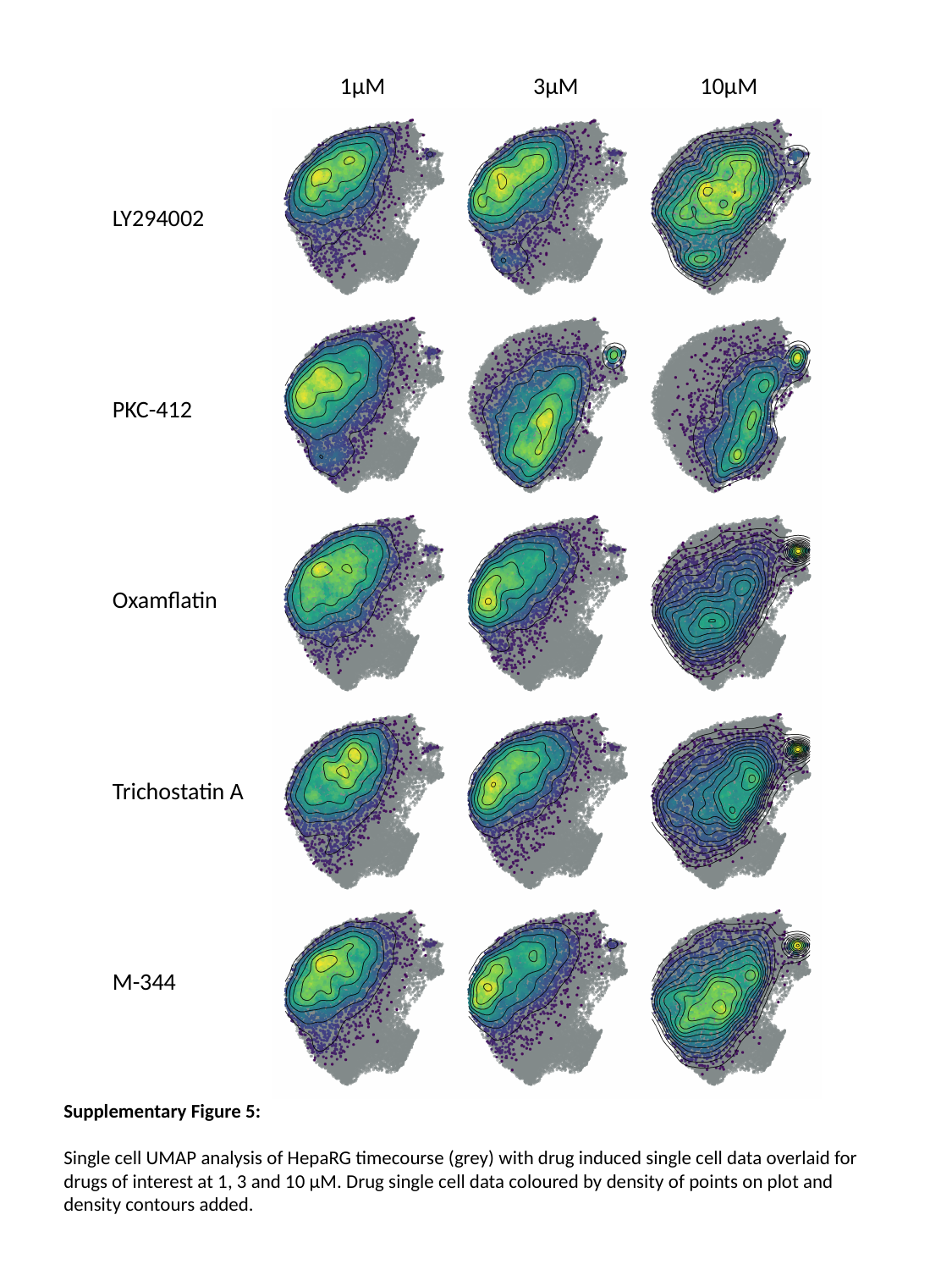

| 1µM 3µM 10µM |
| --- |
| LY294002 |
| PKC-412 |
| Oxamflatin |
| Trichostatin A |
| M-344 |
Supplementary Figure 5:
Single cell UMAP analysis of HepaRG timecourse (grey) with drug induced single cell data overlaid for drugs of interest at 1, 3 and 10 µM. Drug single cell data coloured by density of points on plot and density contours added.

### Slide 6
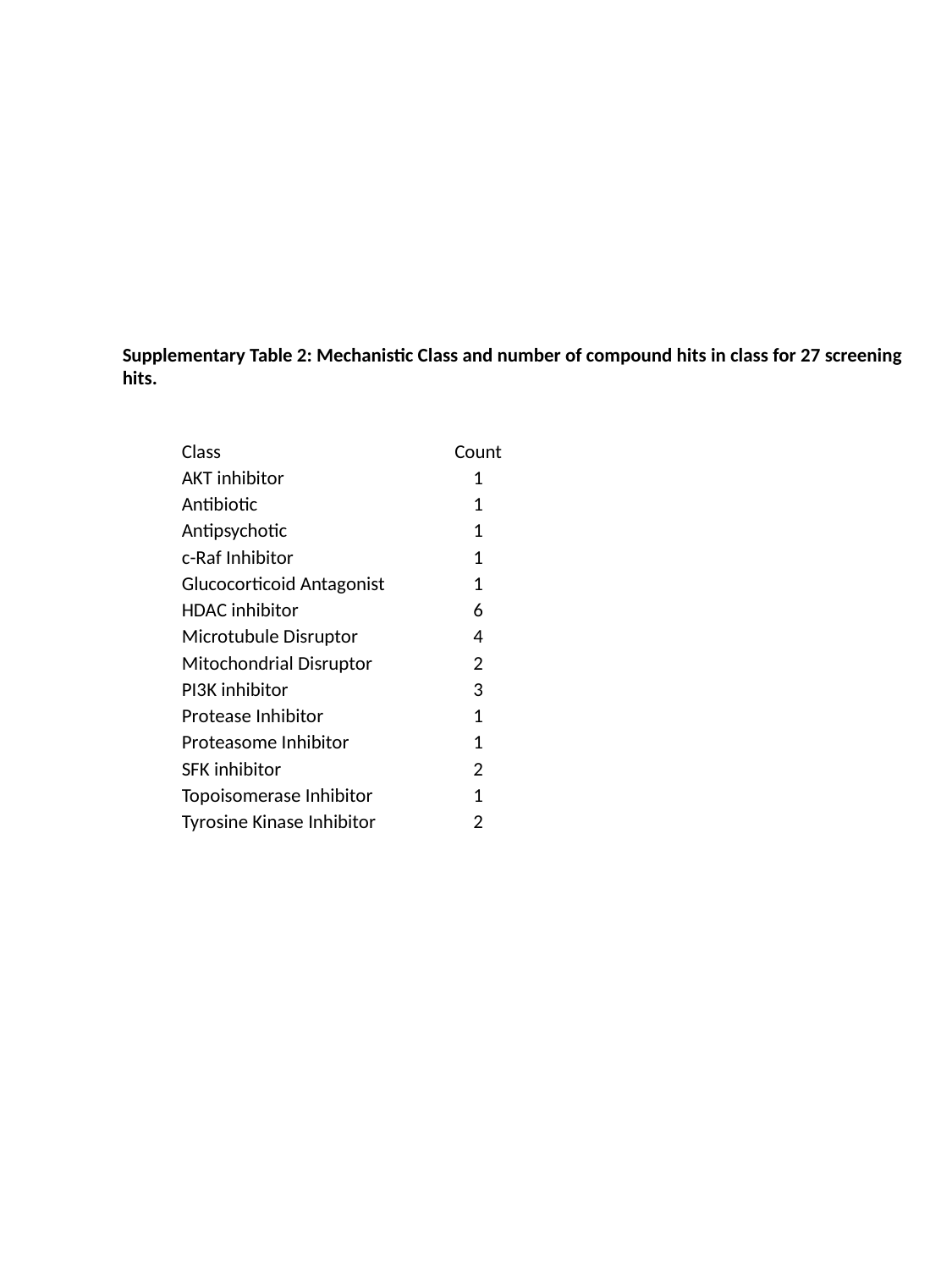

#
Supplementary Table 2: Mechanistic Class and number of compound hits in class for 27 screening hits.
| Class | Count |
| --- | --- |
| AKT inhibitor | 1 |
| Antibiotic | 1 |
| Antipsychotic | 1 |
| c-Raf Inhibitor | 1 |
| Glucocorticoid Antagonist | 1 |
| HDAC inhibitor | 6 |
| Microtubule Disruptor | 4 |
| Mitochondrial Disruptor | 2 |
| PI3K inhibitor | 3 |
| Protease Inhibitor | 1 |
| Proteasome Inhibitor | 1 |
| SFK inhibitor | 2 |
| Topoisomerase Inhibitor | 1 |
| Tyrosine Kinase Inhibitor | 2 |

### Slide 7
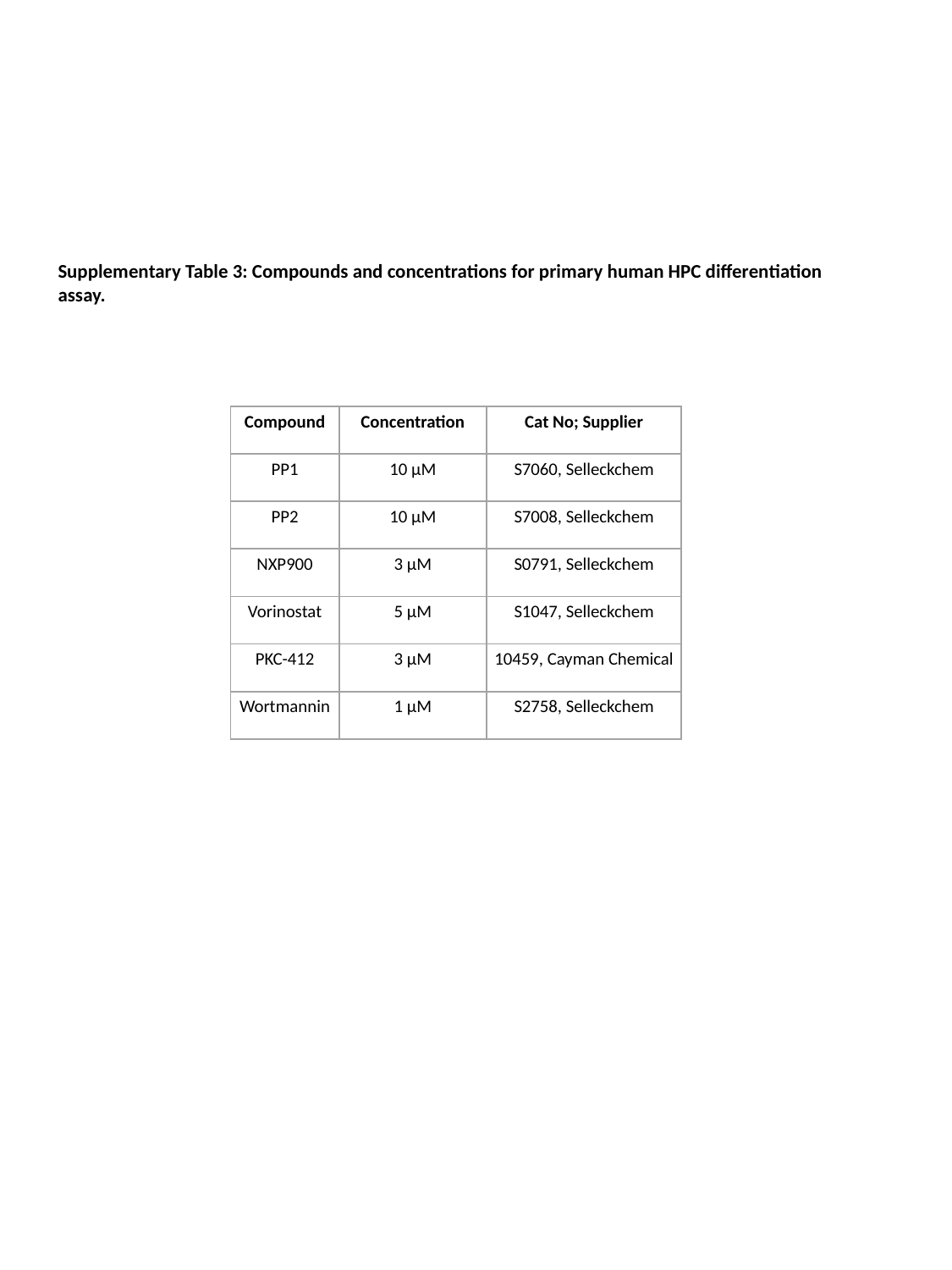

Supplementary Table 3: Compounds and concentrations for primary human HPC differentiation assay.
| Compound | Concentration | Cat No; Supplier |
| --- | --- | --- |
| PP1 | 10 μM | S7060, Selleckchem |
| PP2 | 10 μM | S7008, Selleckchem |
| NXP900 | 3 μM | S0791, Selleckchem |
| Vorinostat | 5 μM | S1047, Selleckchem |
| PKC-412 | 3 μM | 10459, Cayman Chemical |
| Wortmannin | 1 μM | S2758, Selleckchem |
